# Non-literal language processing is jointly supported by the language and Theory of Mind networks: Evidence from a novel meta-analytic fMRI approach

**DOI:** 10.1101/2022.03.08.481056

**Authors:** Miriam Hauptman, Idan Blank, Evelina Fedorenko

## Abstract

Going beyond the literal meaning of utterances is key to communicative success. However, the mechanisms that support non-literal inferences remain debated. Using a novel meta-analytic approach, we evaluate the contribution of linguistic, social-cognitive, and executive mechanisms to non-literal interpretation. We identified 74 fMRI experiments (n=1,430 participants) from 2001-2021 that contrasted non-literal language comprehension with a literal control condition, spanning ten phenomena (e.g., metaphor, irony, indirect speech). Applying the activation likelihood estimation approach to the 825 activation peaks yielded six left-lateralized clusters. We then evaluated the locations of both the individual-study peaks and the clusters against probabilistic functional atlases (cf. macroanatomy, as is typically done) for three candidate brain networks—the language-selective network (Fedorenko et al., 2011), which supports language processing, the Theory of Mind (ToM) network (Saxe & Kanwisher, 2003), which supports social inferences, and the domain-general Multiple-Demand (MD) network (Duncan, 2010), which supports executive control. These atlases were created by overlaying individual activation maps of participants who performed robust and extensively validated ‘localizer’ tasks that target each network in question (n=806 for language; n=198 for ToM; n=691 for MD). We found that both the individual-study peaks and the ALE clusters fell primarily within the language network and the ToM network. These results suggest that non-literal processing is supported by both i) mechanisms that process literal linguistic meaning, and ii) mechanisms that support general social inference. They thus undermine a strong divide between literal and non-literal aspects of language and challenge the claim that non-literal processing requires additional executive resources.

## Introduction

Communicative success often requires going beyond the literal meaning of utterances (e.g., Grice, 1975; Sperber & Wilson, 1986). Neurotypical adults engage in such ‘non-literal’ interpretation—from understanding metaphors, to interpreting sarcasm, to using prosodic cues— seemingly effortlessly. However, the cognitive and neural mechanisms that enable comprehension of non-literal language remain debated.

Early patient investigations linked difficulties in non-literal interpretation with damage to the *right hemisphere (RH)* (e.g., Winner & Gardner, 1977; Myers & Linebaugh, 1981; Delis et al., 1983; Brownell et al., 1983; 1986; Bryan, 1988; Joanette et al., 1989; Weylman et al., 1989; Burgess & Charello, 1996; Myers, 1998). More recently, a growing number of empirical studies have begun to challenge the idea of RH dominance in non-literal processing: brain imaging experiments (e.g., Rapp et al., 2004; Lee & Dapretto, 2006; Bosco et al., 2017; see also Oliveri et al., 2004), meta-analyses of such studies (e.g., Bohrn et al., 2012; Rapp et al., 2012; Reyes-Aguilar et al., 2018), and patient investigations (Ianni et al., 2014; Cardillo et al., 2018; Klooster et al., 2020; see Giora et al., 2000; Zaidel et al., 2002; Klepousniotou & Baum, 2005 for evidence of bilateral involvement) have instead implicated the *left hemisphere (LH)*.

Furthermore, both the left and the right hemisphere each contain multiple distinct functional networks, including the language-selective network (e.g., Fedorenko et al., 2011) and its right homotope, the Theory of Mind network (e.g., Saxe & Kanwisher, 2003), and the domain-general Multiple Demand network (e.g., Duncan, 2010). These networks are associated with distinct cognitive operations, all of which have been argued to contribute to non-literal language comprehension, including in individuals with communication disorders: linguistic processing (e.g., Papagno & Genoni, 2004; Beaty & Silvia, 2013; Whyte & Nelson, 2015), social inference (e.g., Sperber & Wilson, 1986; Happé, 1993; Winner et al., 1998), and executive control (e.g., McDonald & Pearce, 1998; Champagne-Lavau & Stip, 2010).

In recent years, the field of cognitive neuroscience has begun to move away from traditional group analyses, in which brain activations are averaged voxel-wise across individuals and the activation peaks are interpreted via reverse inference from anatomy to function (Poldrack, 2006, 2011; Fedorenko, 2021). Given evidence that functional areas vary in their precise locations in individual brains (e.g., Fischl et al., 2008; Frost & Goebel, 2011; Tahmasebi et al., 2012), the use of individual-subject-level analyses, or ‘precision fMRI’, has become increasingly widespread. In this type of analysis, the relevant functional areas and networks are identified in individual participants using ‘localizer’ tasks that target particular functional areas/networks (e.g., Kanwisher et al., 1997; Saxe et al., 2006; Fedorenko et al., 2010, 2013; Shashidara et al., 2019; Fedorenko, 2021; Gratton & Braga, 2021). Although the shift away from traditional group analyses promises to address questions that could not be answered when the field was dominated by this approach (e.g., Fedorenko et al., 2012; Deen et al., 2015; Braga & Buckner, 2017), ideally, we would not simply abandon many hundreds of past group-level fMRI studies. However, an analysis method that integrates findings from both individual-subject functional localization and traditional group-averaging experiments has not yet emerged.

Here, we present a novel meta-analytic approach for interpreting past fMRI studies and apply it to non-literal language processing. In particular, we leverage *probabilistic functional atlases* (e.g., Dworetsky et al., 2021; Thirion et al., 2021; Lipkin et al., in press), which are created from large numbers of individuals (n>100 for all atlases) who have performed extensively validated functional localizers that selectively target particular functional networks. Given that these atlases capture inter-individual variability in the locations of the relevant networks, we can estimate for any given location in the common space the probability that it belongs to each candidate network. In other words, the probabilistic atlases provide a layer of information beyond anatomy. For example, a particular location in the common space may have a non-zero probability of belonging to two adjacent networks, and by comparing the relative probabilities associated with each network, we may infer which network is more likely to include this location.

This new approach differs from other commonly used meta-analytic approaches, particularly the Activation Likelihood Estimation (ALE; Turkeltaub et al., 2002; Eickhoff et al., 2009) approach. The interpretation of ALE estimates still relies on anatomical landmarks, or on comparisons between the results of multiple meta-analyses performed on group-level data. As such, inter-individual variability in functional architecture is not taken into account. This means that the functional resolution of ALE—its ability to discriminate between nearby functional networks—is low (the logic described in Nieto-Castañón & Fedorenko, 2012 for simple group analyses also applies to ALE meta-analyses). The current approach also differs from NeuroSynth (Yarkoni et al., 2011), as NeuroSynth involves comparing brain areas that are associated with subjectively selected keywords. Because probabilistic atlases are constructed using tasks that have been selectively and robustly linked to particular cognitive processes, relating peak locations to these atlases affords a higher degree of interpretability and a straightforward way to link the results to those from studies that rely on functional localization.

In the present study, we examined data from past fMRI studies (74 studies, 825 activation peaks) that contrasted neural responses to non-literal vs. literal conditions across diverse phenomena with respect to three candidate networks: the language-selective network (Fedorenko et al., 2011), which supports language processing, the Theory of Mind (ToM) network (Saxe & Kanwisher, 2003), which supports social inferences, and the domain-general Multiple Demand (MD) network (Duncan, 2010), which supports executive control. In addition to incorporating probabilistic functional atlases in a novel manner, our meta-analysis improves upon past meta-analyses of non-literal language (e.g., Bohrn et al., 2012; Rapp et al., 2012; Reyes-Aguilar et al., 2018) in that we a) include a larger number of studies, b) focus on targeted, more interpretable contrasts (i.e., non-literal > literal; cf. non-literal > fixation), and c) examine a larger number of non-literal phenomena. To foreshadow our results, we find support for the role of the language-selective and ToM networks, but not the MD network, in non-literal interpretation. Further, in line with past meta-analyses (Bohrn et al., 2012; Rapp et al., 2012; Reyes-Aguilar et al., 2018), we do not find support for an RH bias: the majority of peaks fall in the LH.

## Materials and Methods

### Article selection criteria

The literature search was conducted in accordance with PRISMA guidelines (Moher et al., 2009). Relevant studies were identified in *NeuroSynth*, *Google Scholar*, *APA PsycInfo*, and *PubMed* databases using Boolean searches of the following keywords: “anaphora,” “anthropomorphism,” “comedy,” “discourse comprehension,” “figurative language,” “figure of speech,” “hyperbole,” “humor”, “idioms,” “indirect request,” “indirect speech,” “ironic,” “irony,” “jokes,” “lying,” “metaphor,” “metonymy,” “narrative,” “non-literal language,” “oxymoron,” “paradox,” “personification,” “platitude,” “pragmatics,” “prosody,” “proverbs,” “pun,” “sarcasm,” “sarcastic,” “saying,” “speech act,” “synecdoche,” “text coherence,” “text comprehension,” “understatement”, “fMRI,” “brain,” and “neuroimaging.” Note that we defined “non-literal language” as encompassing any linguistic input whose meaning cannot be reduced to the literal meanings of its parts. As a result, this definition both includes paradigmatic phenomena like metaphors and idioms and extends to phenomena such as establishing text coherence/constructing broader thematic meaning for a narrative text (e.g., Johnson-Laird, 1983; Mason & Just, 2009), as well as prosody.

We additionally examined the reference lists of past neuroimaging meta-analyses on non-literal language processing to minimize the possibility of missing relevant studies (Ferstl et al., 2008; Bohrn et al., 2012; Rapp et al., 2012; Vartanian, 2012; Vrticka et al., 2013; Lisofsky et al., 2014; Yang, 2014; Yang & Shu, 2016; Reyes-Aguilar et al., 2018; Farkas et al., 2021). (Note that we had originally planned to use the NeuroSynth database of extracted peaks (Yarkoni et al., 2011); however, we discovered that NeuroSynth automatically extracts peaks from SPM-style tables in published papers, and such tables frequently include peaks for contrasts in both directions: e.g., in our case, non-literal>literal and literal>non-literal. As a result, we opted to manually extract peaks from only non-literal>literal contrasts.)

260 articles were selected for full-text screening based on the content of their abstracts. The selection criteria included: (1) fMRI (not PET or MEG/EEG) was used; (2) participants were neurotypical, and not aging, adults; (3) participants were native speakers of the language in which the experiment was conducted; (4) a standard whole-brain random effects analysis (Holmes & Friston, 1998) was performed; (5) activation peaks were reported in Talairach (Talairach & Tournoux, 1988) or Montreal Neurological Institute (MNI) (Evans et al., 1993) coordinate systems; and (6) contrasts targeted non-literal language comprehension, in the listening or reading modality, versus a (literal) linguistic baseline (cf. coarser-grain contrasts like non-literal language processing vs. fixation). The 74 studies that satisfied these criteria were published between 2001 and 2021 and targeted ten linguistic phenomena (**Table 1**). Across these 74 studies, 825 activation peaks (from 102 contrasts, between 1 and 4 contrasts per study) were extracted for analysis. Participants included 1,430 individuals (between 8 and 39 per study; *M* = 19.2) aged 18 to 55 (*M* = 24.8 years), 55% female (see **SI Table 1** for further details). **Table 1** summarizes the distribution of studies and peaks across the ten phenomena.

**Table 1.** Distribution of studies and peaks across linguistic phenomena. (One of the 74 studies included separate contrasts for non-sarcastic irony and sarcasm and is therefore counted twice in the table.) Studies on prosody that focused on the role of emotionally charged prosodic cues were excluded.

| Linguistic phenomenon | Number of studies | Number of peaks |
| --- | --- | --- |
| Humor | 8 | 121 |
| Idiom | 6 | 54 |
| Indirect Speech | 10 | 160 |
| Irony | 10 | 68 |
| Metaphor | 19 | 201 |
| Metonymy | 2 | 10 |
| Prosody | 4 | 64 |
| Proverb | 1 | 9 |
| Sarcasm | 2 | 10 |
| Text Coherence | 13 | 128 |
| **TOTAL** | **74 unique studies** | **825** |

### The probabilistic functional atlases for the three brain networks of interest

The individual activation peaks and the clusters derived from these peaks via the standard activation likelihood estimation (ALE) analysis (e.g., Turkeltaub et al., 2002; Eickhoff et al., 2009), as described below (section <u>Activation likelihood estimation (ALE) analysis</u>), were evaluated with respect to probabilistic functional atlases for three candidate networks of interest: the language network, the Theory of Mind (ToM) network, and the Multiple Demand (MD) network. For each network, an activation overlap map was created by overlaying a large (n>100) number of individual binarized activation maps for the ‘localizer’ task targeting each network, as described below (this is the first step in the Group-Constrained Subject-Specific analytic approach, as described in Fedorenko et al., 2010 and Julian et al., 2012). To account for inter-individual variability in the overall level of activation, we selected in each individual the top 10% most localizer-responsive voxels across the brain (fixed-statistical-threshold approaches yield near-identical results; Lipkin et al., 2022). Specifically, we sorted the *t*-values for the relevant contrast and took 10% of voxels per participant with the highest values. In the resulting activation overlap map, the value in each voxel represents the number of participants for whom that voxel belongs to the top 10% most localizer-responsive voxels. These values—turned into proportions by dividing each value by the total number of participants that contribute to the overlap map—can then be used to estimate the probability that a given voxel belongs to the target network. Consider the extreme cases: if a voxel does not belong to the top 10% most localizer-responsive voxels in *any* participant, that voxel is extremely unlikely to belong to the network of interest, and if a voxel belongs to the top 10% most localizer-responsive voxels in *every* participant, that voxel is extremely likely to belong to the network of interest. In practice, for brain networks that support high-level cognitive functions and fall within the association cortex, network probability is unlikely to ever be 1 because of high inter-individual variability in the precise locations of these networks (e.g., Frost & Goebel, 2011; Tahmasebi et al., 2012), as discussed above. Our degree of confidence in assigning a voxel to a network will therefore be constrained by the maximum inter-subject overlap value for that network.

The task used to localize the ***language network*** is described in detail in Fedorenko et al. (2010) and targets brain regions that support high-level language processing, including phonological, lexical-semantic, and combinatorial (semantic and syntactic) processes (e.g., Fedorenko et al., 2010, 2012, 2016, 2020; Bautista & Wilson, 2016; Blank et al., 2016; Regev et al., 2021). It also identifies right-hemisphere homotopes of the left-hemisphere language regions (e.g., Mahowald & Fedorenko, 2016), which have been proposed to play a role in non-literal language comprehension / pragmatic reasoning (e.g., Joanette et al.,1990; Kuperberg et al., 2000; Mashal et al., 2005; Coulson & Williams, 2005; Diaz & Hogstrom, 2011; Eviatar & Just, 2006). Briefly, we used a reading task that contrasted sentences (the critical condition) and lists of unconnected, pronounceable nonwords (the control condition; **Figure 1**) in a standard blocked design with a counterbalanced condition order across runs. By design, this localizer contrast subtracts out lower-level perceptual (speech or reading-related) and articulatory motor processes (see Fedorenko & Thompson-Schill, 2014 for discussion) and has been shown to generalize across materials, tasks, visual/auditory presentation modality, and languages (e.g., Fedorenko et al., 2010; Fedorenko, 2014; Scott et al., 2017; Ivanova et al., 2020; Chen et al., 2021; Malik-Moraleda, Ayyash et al., 2021). Further, this network emerges robustly from task-free naturalistic data (e.g., Braga et al., 2020; see also Blank et al., 2014, Paunov et al., 2018). Participants read the stimuli one word/nonword at a time in a blocked design, with condition order counterbalanced across runs. Each sentence/nonword sequence was followed by a button-press task to maintain alertness. A version of this localizer is available from https://evlab.mit.edu/funcloc/download-paradigm, and the details of the procedure and timing are described in **Figure 1** and **Table 2**. The probabilistic functional atlas used in the current study (Language Atlas (LanA); Lipkin et al., 2022) was constructed using data from 806 participants, and the voxel with the highest network probability had a value of 0.82 (i.e., belonged to the top 10% of most language-responsive voxels in 82% of participants).

**Fig. 1.**
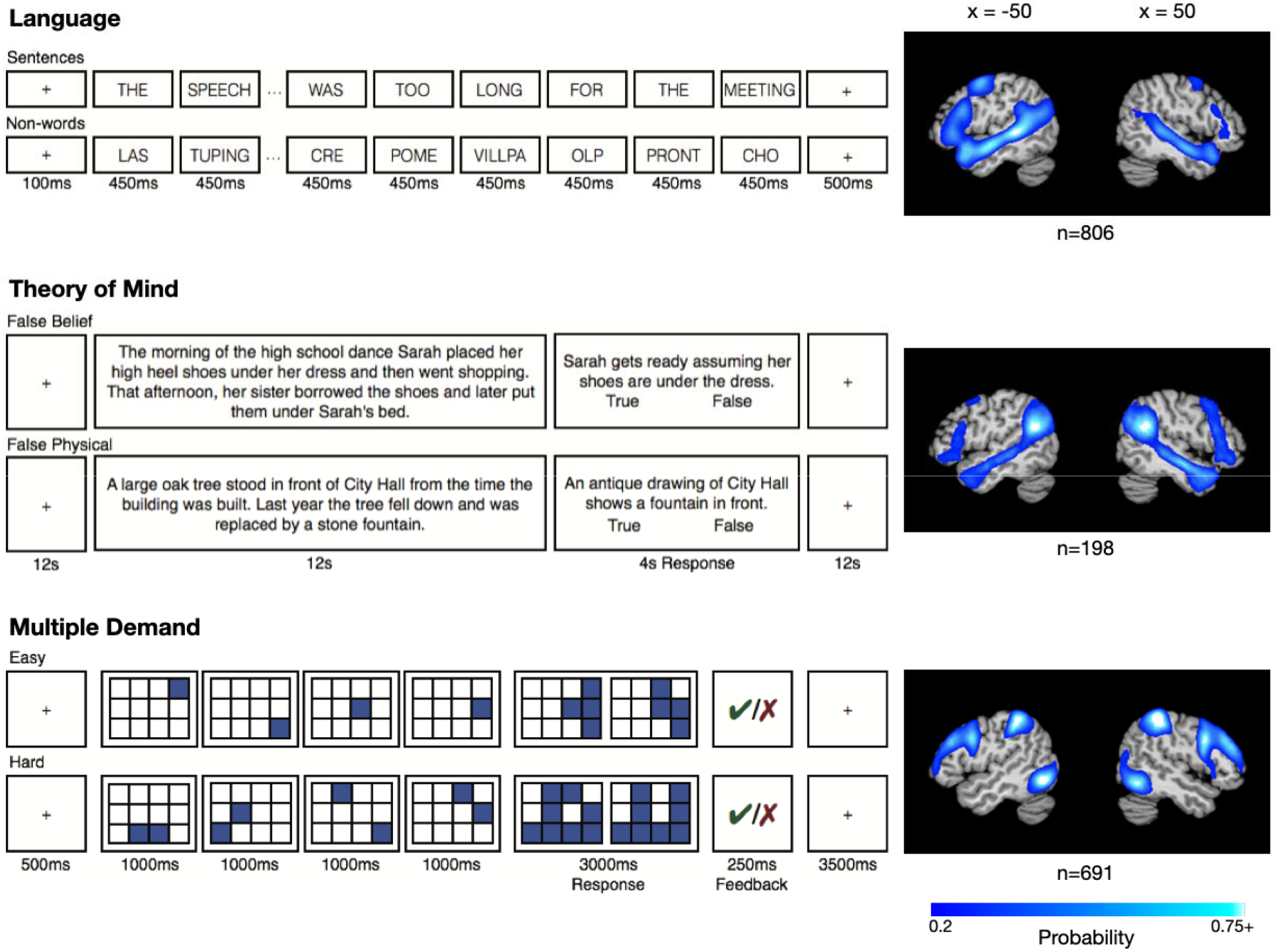
The three functional localizer paradigms (language, ToM, MD) and the resulting probabilistic functional atlases. The maps illustrate, for each voxel, the proportion of participants for whom that voxel belongs to the top 10% of localizer-responsive voxels.

**Table 2.** Details on the design and timing of the language, ToM and MD localizer tasks.

|  | **Language** | **ToM** | **MD** |
| --- | --- | --- | --- |
| **Design** | Blocked | Long-event-related | Blocked |
| **Length of trial** | 4.8-18 s | 14 s | 8-9 s |
| **Trials per block** | 1-5 | N/A | 2-4 |
| **Blocks/events per run** | 12-18 | 10 | 10-12 |
| **Blocks/events per condition per run** | 4-16 | 5 | 5-6 |
| **Length of run** | 336-504 s | 272 s | 288-448 s |
| **Runs** | 2-8 | 1-2 | 2-4 |
| **Versions of the design** | 10 | 1 | 3 |

The task used to localize the ***ToM network*** is described in detail in Saxe and Kanwisher (2003) and targets brain regions that support reasoning about others’ mental states. Briefly, we used a task based on the classic false belief paradigm (Wimmer & Perner, 1983) that contrasted verbal vignettes about false beliefs (e.g., a protagonist has a false belief about an object’s location; the critical condition) versus linguistically matched vignettes about false physical states (physical representations depicting outdated scenes, e.g., a photograph showing an object that has since been removed; the control condition; **Figure 1**). By design, this localizer focuses on ToM reasoning to the exclusion of affective or non-propositional aspects of mentalizing and has been shown to generalize across materials (verbal and non-verbal), tasks, and visual/auditory presentation modality (e.g., Gallagher et al., 2000; Saxe & Kanwisher, 2003; Saxe et al., 2006; Jacoby et al., 2016). Furthermore, this network emerges robustly from task-free naturalistic data (e.g., Braga & Buckner, 2017; DiNicola et al., 2020; see also Paunov et al., 2018). Participants read the vignettes one at a time in a long-event-related design, with condition order counterbalanced across runs. Each vignette was followed by a true/false comprehension question. A version of this localizer is available from http://saxelab.mit.edu/use-our-efficient-false-belief-localizer, and the details of the procedure and timing are described in **Figure 1** and **Table 2**. The probabilistic functional atlas used in the current study (unpublished data from the Fedorenko lab) was constructed using data from 198 participants, and the voxel with the highest network probability had a value of 0.88.

The task used to localize the ***MD network*** is described in detail in Fedorenko et al. (2013) (see also Blank et al., 2014) and targets brain regions that are sensitive to general executive demands. Briefly, we used a spatial working memory task that contrasted a harder and an easier condition. On each trial, participants saw a 3 x 4 grid and kept track of eight (the critical, harder, condition) or four (the control, easier, condition; **Figure 1**) locations that were sequentially flashed two at a time or one at a time, respectively. Participants indicated their memory for these locations in a two-alternative, forced-choice paradigm via a button press. Feedback was provided after every trial. There is ample evidence that demanding tasks of many kinds activate this network (e.g., Duncan & Owen 2000; Fedorenko et al. 2013; Hugdahl et al. 2015; Shashidara et al., 2019; and Assem et al., 2020a). Furthermore, this network emerges robustly from task-free naturalistic data (e.g., Assem et al., 2020a; Braga et al., 2020; see also Blank et al., 2014, Paunov et al., 2019). Hard and easy conditions were presented in a blocked design, with condition order counterbalanced across runs. This localizer is available from the authors upon request and the details of the procedure and timing are described in **Figure 1** and **Table 2**. The probabilistic functional atlas used in the current study (MDAtlas; Lipkin et al., in prep.) was constructed using data from 691 participants, and the voxel with the highest network probability had a value of 0.75. (It is worth noting that the three networks of interest differ somewhat in the range of their non-zero network probability values: language = 0.001-0.82, ToM = 0.005-0.88, and MD = 0.001-0.75. We decided against normalizing these values for each map (so that the highest network probability would be set to 1, or to the highest value observed across any of the three networks) because doing so would obscure meaningful differences in the likelihood that a given voxel belongs to each network.)

Critically, these three networks are robustly spatially and functionally dissociable within individuals in both naturalistic (e.g., Blank et al., 2014; Paunov et al., 2018; Braga et al., 2020) and task-based fMRI paradigms despite their close proximity to each other within the association cortex. The ***language network*** selectively supports linguistic processing, showing little or no response to diverse executive function tasks (e.g., Fedorenko et al., 2011; Monti et al., 2012; see Fedorenko & Blank, 2020 for a review) and mentalizing tasks (Deen et al., 2015, Paunov, 2018; Paunov et al., 2021; Shain, Paunov, Chen et al., 2022). The data from the patient literature mirrors this selectivity: damage to the language network does not appear to lead to difficulties in executive or social processing (see Fedorenko and Varley, 2016 for a review). The ***ToM network*** selectively supports social cognition, showing little or no response to linguistic input without mental state content (Deen et al., 2015; Paunov et al., 2021; Deen & Freiwald, 2021; Shain, Paunov, Chen et al., 2022) or to executive demands (Saxe et al., 2006; Scholz et al., 2009; Willems et al., 2010). Finally, the ***MD network*** supports diverse executive demands (e.g., Duncan, 2010, 2013; Assem et al., 2020a; Smith et al., 2021) and is linked to fluid reasoning ability (e.g., Woolgar et al., 2010; Assem et al., 2020b), but plays a limited role in language comprehension once task demands are controlled for (e.g., Diachek, Blank, Siegelman et al., 2020; Shain, Blank et al., 2020; Wehbe et al., 2021; see Fedorenko & Shain, 2021 for a review), and in social cognition (e.g., Willems et al., 2010). Because of the selective relationship between each of the three networks and a particular set of cognitive processes, activity in these networks can be used as an index of the engagement of the relevant processes (see e.g., Mather et al., 2013, for discussion), circumventing the need for precarious reverse inference from anatomical locations to function (e.g., Poldrack et al., 2006, 2011; Fedorenko, 2021). This is the approach that is adopted in studies that rely on functional localization (e.g., Brett et al., 2002; Saxe et al., 2006; Fedorenko, 2021). Here, we extend this general logic to the meta-analysis of group-level activation peaks by leveraging information about the landscape of each network of interest based on probabilistic functional atlases for the relevant localizer tasks.

### Extracting network probabilities for the individual-study peaks

Prior to analysis, activation peaks that were reported in Talairach space were converted to the MNI space using icbm2tal transform SPM conversion in GingerALE 3.0.2 (Eickhoff et al., 2009, 2012). The MNI coordinates of the 825 activation peaks included in the dataset were tested against the probabilistic functional atlases (created as described above, section <u>The probabilistic functional atlases for the three brain networks of interest</u>) for each of the three functional networks of interest. For each coordinate, three network probability values were extracted (one from each functional atlas). (Note that because of the variability in the precise locations of functional areas and the proximity of the three networks to each other in parts of the association cortex, many voxels have non-zero values for more than a single network (despite the fact that there is little to no overlap among the networks in individual participants)—this is precisely the argument against traditional group-averaging analyses in fMRI; e.g., Fedorenko & Blank, 2020; DiNicola & Buckner, 2021; Fedorenko, 2021; Gordon & Nelson, 2021; Gratton & Braga, 2021; Smith et al., 2021).

### Activation likelihood estimation (ALE) analysis

In addition to examining the individual-study activation peaks, we used the GingerALE software (Eickhoff et al., 2009, 2012) to perform a traditional fMRI meta-analysis via the activation likelihood estimation (ALE) method (Turkeltaub et al., 2002). ALE identifies regions that show consistent activation across experiments (Eickhoff et al., 2009, 2012). The clusters that ALE yields should be less noisy than the individual-study activation peaks, especially when multiple-comparison correction is not appropriately applied and participant sample sizes are small in individual studies (e.g., Genovese et al., 2002; Eklund et al., 2016; Chen et al., 2018).

The 74 studies in our dataset yielded 825 activation peaks (**Table 1**; see **SI Table 1** for a complete list of studies), but 39 peaks (fewer than 5%) fell outside the MNI template used by GingerALE and were therefore excluded, leaving 786 peaks. For each study, a map was created in which each voxel in the MNI space received a modeled activation score. Modeled activation scores reflect the likelihood that significant activation for one particular experiment was observed at a given voxel. The 74 modeled activation maps were then unified, generating an ALE value for each voxel.

Significance was assessed by comparing the observed ALE values to a null distribution that was generated by repeatedly calculating ALE values using randomly placed activation peaks (1,000 permutations). A cluster-forming threshold of p<0.001 uncorrected identified contiguous volumes of significant voxels (“clusters”), and clusters that survived a cluster-level family-wise error (FWE) of p<0.05 were considered significant. Cluster-level FWE correction has been argued to be the most appropriate correction for ALE, as it minimizes false positives while remaining more sensitive to true effects in comparison to other correction methods (Eickhoff et al., 2016).

Applying ALE to the 74 experiments yielded six significant clusters that were located in the left frontal and temporal cortex, and in the left amygdala (**SI Figure 1**). The clusters varied in size from 176 to 1,695 voxels.

### Critical analyses

We asked two key research questions: the first, critical question examines the locations of the activation peaks with respect to the *three brain networks of interest*: the language network, the ToM network, and the MD network, and the second question is motivated by the prior claim about the privileged role of the *right hemisphere* in non-literal language processing (e.g., Winner & Gardner, 1977). Note that in all the analyses, we collapse the data from across the ten phenomena. We adopted this approach because i) none of the phenomena have a sufficient number of studies to enable meaningful phenomenon-level examination, and ii) there is currently a lack of consensus about the ways to carve up the space of non-literal phenomena (see <u>Discussion</u>). However, in an exploratory analysis, we examined the contribution of different phenomena to the peaks/clusters that load on the functional networks.

#### Q1: How are the activation peaks from prior fMRI studies of non-literal language comprehension distributed across the language, ToM, and MD networks?

##### Analysis of individual-study peaks

We evaluated the locations of the activation peaks with respect to our three brain networks of interest (language, ToM, MD). Almost all the peaks—820/825 (99%)—were observed in voxels with a non-zero network probability in at least one functional atlas. To test whether the network probabilities associated with the activation peaks differed between the networks, we developed the following statistical procedure (note that for all analyses, we excluded the 13 peaks that fell on the cortical midline, i.e., had an x coordinate value of 0, given that we wanted to examine each network in each hemisphere separately, which left 812 peaks for analysis):

1. For each of the 812 peaks, we calculated the difference in network probabilities between each pair of networks (language vs. ToM, language vs. MD, and ToM vs. MD), and recorded the median (across peaks) of these difference values for each of the three contrasts in each hemisphere separately. (We used median instead of mean values to better capture the central tendencies given the skewness of the distributions.)
2. We then created 3D maps from which random peak sets could be selected (against which the location of true observed peaks could be evaluated, as described in Step 4 below). Specifically, we created a map for each hemisphere whereby we removed the voxels whose locations were associated with a network probability of 0 in all three functional atlases (for the language, ToM, and MD networks). This restriction was imposed so as to a) constrain the locations of the baseline voxel sets to the parts of the brain where true observed peaks were found (as noted above, almost all the peaks were observed in voxels that had a non-zero network probability in at least one network); and thus, b) to construct a more conservative test, making it more difficult to detect between-network differences.
3. In each hemisphere, the same number of peaks as in our dataset (n=490 in the LH, and n=322 in the RH) were then randomly sampled from the maps that were created in Step 2, and the median difference (across peaks) for each contrast (language vs. ToM, language vs. MD, ToM vs. MD) was computed, as in Step 1. This procedure was repeated 10,000 times, yielding an empirical null distribution of median network probability differences.
4. Finally, we compared the true median network probability differences against the distribution of network probability differences obtained in Step 3 to yield a significance value for each of the three contrasts within each hemisphere.

##### Analysis of activation likelihood estimation (ALE) clusters

We evaluated the locations of 1) the center peaks of the six significant ALE clusters, and 2) all voxels contained within each cluster with respect to our three brain networks of interest (language, ToM, MD). Importantly, whereas individual study peaks might be skewed by a single study with a particularly large number of activation peaks, ALE estimates incorporate the sample size of each study and are therefore less susceptible to this source of bias. To test whether the network probabilities differed between the networks, we evaluated the network probabilities of the voxels in each cluster and, for each cluster individually, subsequently performed the statistical procedure described above (in <u>Analysis of individual-study peaks</u>).

#### Q2: Do the activation peaks from prior fMRI studies of non-literal language comprehension exhibit a right hemisphere bias?

##### Analysis of individual-study peaks

To test whether the two hemispheres differed in the number of the activation peaks, we ran two logistic mixed effect regression models using the “lme4” package in R. The first tested the full set of 812 activation peaks (excluding the 13 peaks that fell on the cortical midline), and the second focused on the subset of the 812 peaks (n=636) that exhibited a network probability of 0.10 or greater in at least one network. The second model was included to ensure that the results are not driven by a subset of peaks that do not load strongly on any of the three networks of interest. For both models, the following formula was used, which included a random intercept for experiment (n=74):

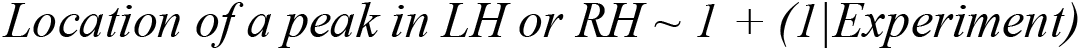

### Analysis of activation likelihood estimation (ALE) clusters

We examined the locations of the six significant ALE clusters with respect to hemisphere.

## Results

### Non-literal language comprehension draws primarily on the language and ToM networks

Across the 812 non-midline peaks, the highest network probability was observed for the ToM network functional atlas (maximum=0.86, mean=0.19, SD=0.18, median=0.12; 802 total peaks with nonzero network probabilities), followed by the language atlas (maximum=0.80, mean=0.17, SD=0.17, median=0.10; 807 nonzero peaks), and the MD atlas (maximum=0.65, mean=0.11, SD=0.14, median=0.04; 806 nonzero peaks). In both hemispheres, the median non-zero network probabilities across the activation peaks numerically exceeded the median non-zero probability values across the functional atlases as a whole: language atlas (LH: peaks=0.12, atlas=0.04; RH: peaks=0.07, atlas=0.04), ToM (LH: peaks=0.13, atlas=0.04; RH: peaks=0.13, atlas=0.04), and MD (LH: peaks=0.03, atlas=0.02; RH: peaks=0.04, atlas=0.02). See **Figure 2** for a depiction of the network probability distributions of the individual-study peaks for each network.

**Fig. 2.**
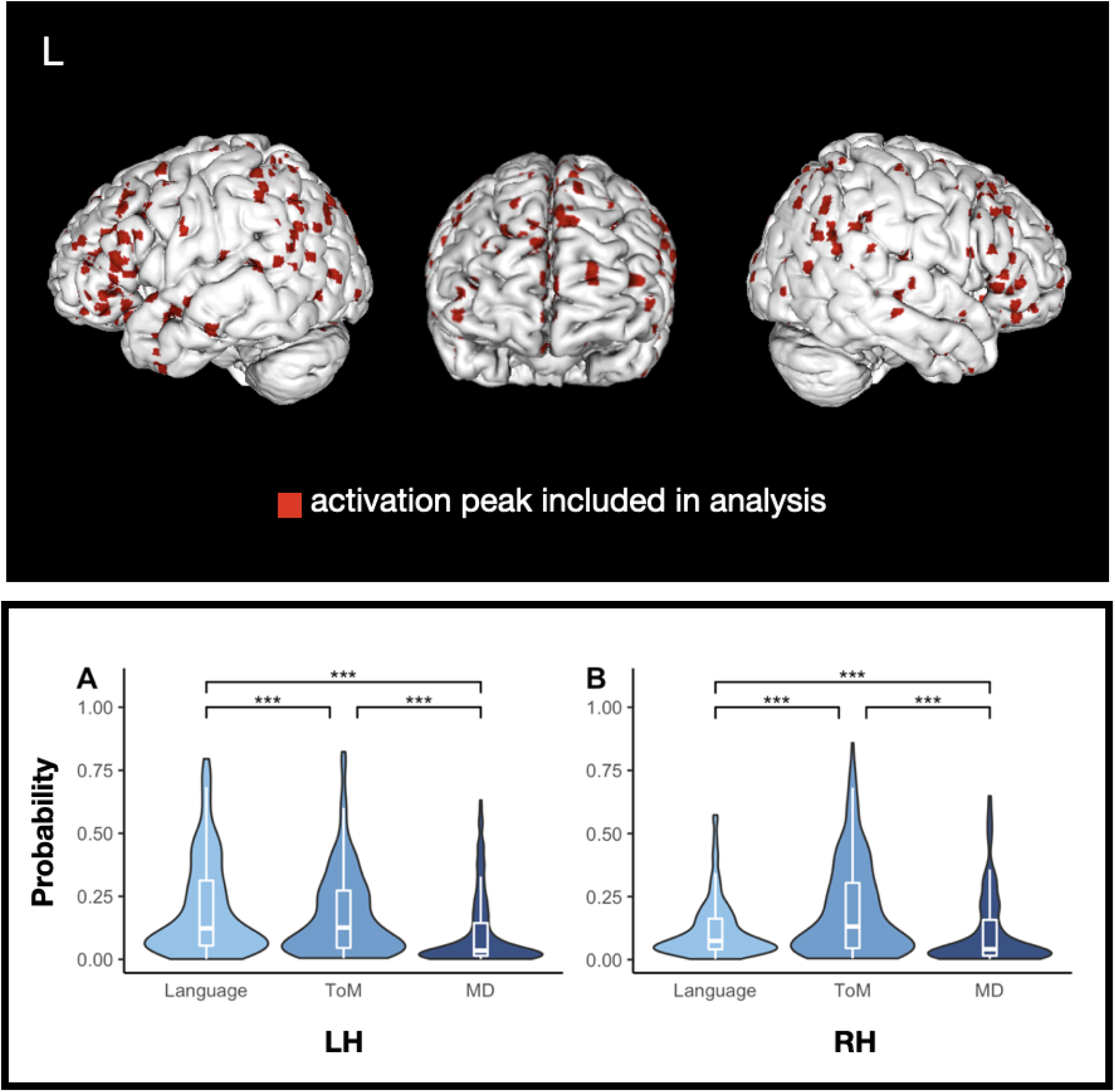
Individual peaks. The top panel displays individual activation peaks plotted on the smoothed (width = 2; kernel size = 3) cortical surface of a high-resolution structural MRI scan in MNI space. The bottom panel displays the distribution of nonzero network probabilities associated with all individual peaks, by hemisphere. Significance of median differences between networks was assessed via a permutation test; all ps < .0001.

We then examined between-network differences in network probabilities. Among the 490 LH peaks, the magnitude of the median difference in network probabilities was highest between the language and MD networks (language vs. MD = 0.07, language vs. ToM = 0.02, ToM vs. MD = 0.04). In the RH (322 peaks), the magnitude of the median difference in network probabilities was highest between the ToM and MD networks (ToM vs. MD = 0.04, language vs. ToM = 0.02, language vs. MD = 0.02). Our permutation analysis (see Methods) revealed that the LH activation peaks were located more centrally in the language network than in either the ToM or the MD network (both ps<0.0001) and more centrally in the ToM network than the MD network (p<0.0001). The RH activation peaks were located more centrally in the ToM network than either the language or the MD network (both ps<0.0001) and more centrally in the language network than the MD network (p<0.0001) (see **Figure 2**). Importantly, this analysis controls for network lateralization (language=LH, ToM=RH) by generating an empirical null distribution of probability differences in each hemisphere separately. Higher network probabilities observed in the language network among the LH voxels and in the ToM network among the RH voxels therefore exceed what would be expected given the (already-skewed) distribution of network probabilities in each hemisphere.

With respect to the ALE clusters (see **Table 3** and **SI Figure 1**), the center peaks of four clusters, including the largest cluster, exhibited the highest network probabilities in the language atlas. Two of these were located in the left temporal lobe (primarily within the superior and middle temporal gyri), one within the left inferior frontal gyrus, and one within the left amygdala (although note that the network probability for the amygdala peak is overall relatively low compared to the cortical peaks). The center peaks of two additional clusters exhibited the highest network probabilities in the ToM atlas. One of these clusters was located in the left medial frontal gyrus and one in the left superior and middle temporal gyri.

**Table 3.** Network probabilities for the six significant ALE clusters. The number of voxels in each cluster is listed under “Size (in voxels)”. The “Broad Anatomical Area” column lists the macroanatomical region that overlaps the most with each cluster (as determined by the ALE). In the “Language,” “ToM,” and “MD” columns, the first value represents the network probability associated with that cluster’s center peak and the second value represents the median network probability of all voxels contained in the cluster. In bold, we highlighted for each cluster the network that has the highest network probabilities.

| Cluster | Size<br>(in<br>voxels) | Hemisphere | Broad<br>Anatomical<br>Area | x | y | z | Language | ToM | MD |
| --- | --- | --- | --- | --- | --- | --- | --- | --- | --- |
| 1 | 283 | LH | STG | -54 | -1 | -16 | **0.743 / 0.634** | 0.348 / 0.298 | 0.001 / 0.009 |
| 2 | 357 | LH | MTG | -55 | -33 | -3 | **0.738 / 0.638** | 0.394 / 0.323 | 0.016 / 0.016 |
| 3 | 1,695 | LH | IFG | -47 | 26 | 3 | **0.424 / 0.298** | 0.273 / 0.177 | 0.030 / 0.064 |
| 4 | 176 | LH | Amygdala | -23 | -10 | -17 | **0.136 / 0.117** | 0.030 / 0.040 | 0.004 / 0.009 |
| 5 | 528 | LH | STG | -49 | -63 | 29 | 0.197 / 0.272 | **0.611 / 0.578** | 0.016 / 0.028 |
| 6 | 454 | LH | MedFG | -3 | 51 | 31 | 0.355 / 0.227 | **0.449 / 0.369** | 0.012 / 0.022 |

Analysis of the network probabilities associated with the full set of voxels comprising each ALE cluster yielded similar results (**Table 3**). The voxels in the four clusters whose center peaks had the highest network probabilities in the language atlas also exhibited the greatest *median network probabilities* in the language atlas, whereas the voxels in the two clusters whose center peaks had the highest network probabilities in the ToM atlas also exhibited the greatest *median network probabilities* in the ToM atlas. Results from the permutation analysis supported this apparent distinction between the four “language” clusters and the two “ToM” clusters: voxels comprising the language clusters had significantly higher network probabilities in the language network than in the ToM and MD networks, and voxels comprising the ToM clusters had significantly higher network probabilities in the ToM network than in the language and MD networks (all ps<0.0001). These results are displayed in **Figure 3**.

**Fig. 3.**
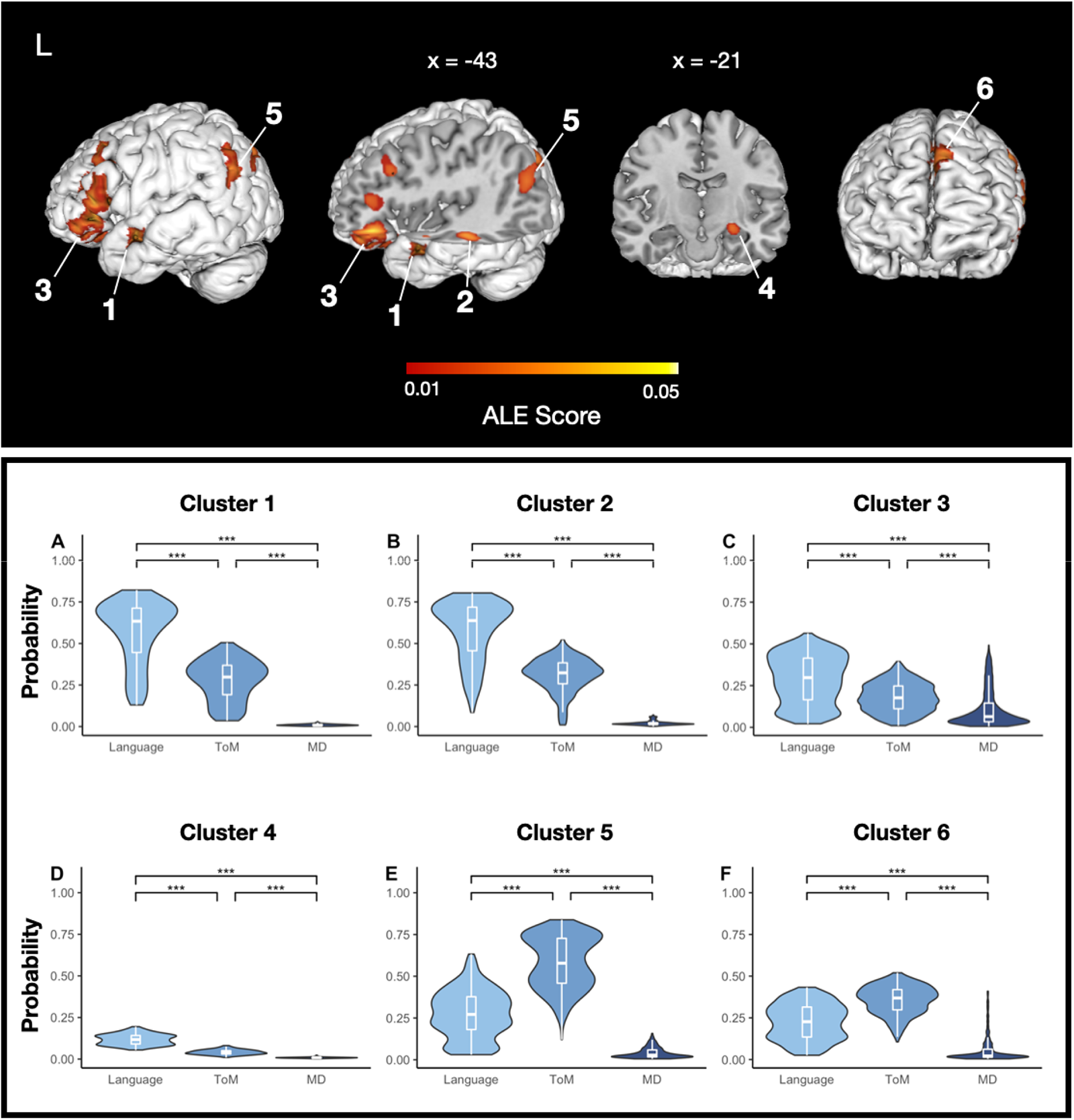
Activation likelihood estimation (ALE) results. The top panel displays ALE scores associated with each of the 6 significant ALE clusters, plotted on the smoothed (width = 2; kernel size = 3) cortical surface of a high-resolution structural MRI scan in MNI space (cluster-forming threshold = p<0.001 uncorrected, cluster-level family-wise error (FWE) = p<0.05, 1000 permutations). The bottom panel displays the distribution of nonzero network probabilities associated with all voxels contained within each cluster. Significance of median differences between networks was assessed via a permutation test; all ps < .0001.

Finally, in an exploratory analysis, we evaluated the phenomena that contributed to the six significant ALE clusters (see **SI Table 2**). No clear differentiation in terms of phenomena that contribute to the language vs. the ToM clusters was apparent.

### No evidence of a RH bias for non-literal language comprehension

Of the full set of 812 peaks (excluding the 13 that fell on the cortical midline), 322 peaks (∼40%) were located in the RH, and 490 (∼60%) were located in the LH (**Figure 4**). This difference was highly reliable (b = -0.44, SE = 0.1, z = -4.42, p<0.001). Similarly, of the subset of the 636 peaks with network probabilities of at least 0.10 in at least one network of interest, 245 peaks (∼39%) were located in the RH, and 391 (∼61%) were located in the LH (**Figure 4**). This difference was highly reliable (b = -0.50, SE = 0.11, z = -4.5, p<0.001). The LH bias was also mirrored in the ALE analysis: all six clusters fell in the LH. These findings provide additional support for the notion that the LH is an important contributor—seemingly more important than the RH—to non-literal language processing (Bohrn et al., 2012; Rapp et al., 2012; Reyes-Aguilar et al., 2018).

**Fig. 4.**
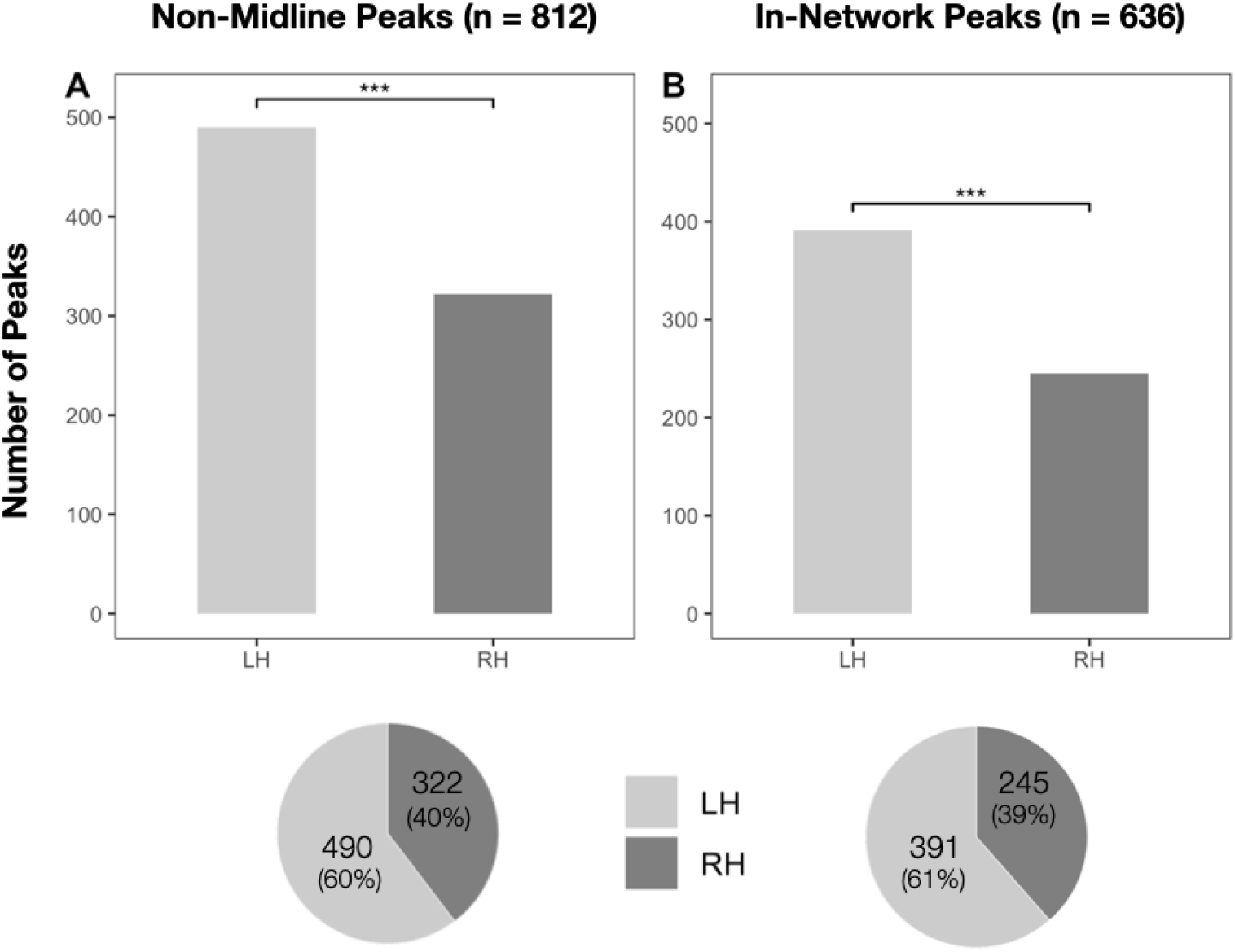
Peaks by hemisphere. In the top panel, A displays the number of non-midline peaks in the left vs. right hemisphere. B displays the left vs. right breakdown of non-midline peaks with an network probability of at least 0.10 in at least one network. Significance was assessed via logistic mixed effects regression; all ps < .001. Pie charts displaying the proportion of activation peaks in the left vs. right hemisphere are shown in the bottom panel.

## Discussion

To illuminate the cognitive and neural bases of non-literal language comprehension, we performed a meta-analysis of group-level activation peaks from past fMRI studies. Specifically, we developed a novel approach that leverages ‘probabilistic functional atlases’ for three brain networks that have been implicated in non-literal comprehension (the language network, the Theory of Mind network, and the Multiple Demand network). The atlases are built using large numbers of individual activation maps for extensively validated ‘localizer’ tasks (e.g., Saxe et al., 2006; Fedorenko, 2021) and provide estimates of the probability that a location in the common brain space belongs to a particular network. Because each of these networks has been rigorously characterized and selectively linked to particular cognitive processes in past work, activation peak locations therein can be interpreted as evidence for the engagement of the relevant process(es) (Mather et al., 2013). This approach is therefore superior to the traditional meta-analytic approach where activation peaks or ALE clusters (Turkeltaub et al., 2002) are interpreted solely based on their anatomical locations or in relation to the results of additional meta-analyses.

The three networks that we examined include i) the language-selective network, which supports literal comprehension, including lexical and combinatorial operations (Fedorenko et al., 2020), ii) the Theory of Mind (ToM) network, which supports social inference, including mentalizing (Saxe & Kanwisher, 2003), and iii) the Multiple Demand (MD) network, which supports executive control (Duncan, 2010). We asked two research questions. The first, critical question concerned the distribution of peaks across the networks. The individual peaks and ALE clusters tended to fall in the language and ToM networks, but not the MD network. The second question, motivated by past patient investigations that have linked non-literal comprehension impairments to RH damage (e.g., Winner & Gardner, 1977), asked whether the peaks were more likely to fall in the RH. In line with past meta-analyses of non-literal language (Bohrn et al., 2012; Rapp et al., 2012; Reyes-Aguilar et al., 2018), we found that the peaks/clusters fell primarily in the LH. Below, we discuss several issues that these results inform, and highlight some outstanding questions and some methodological implications.

### The role of the ToM and language networks in non-literal comprehension

In line with past studies that have reported activations in putative ToM areas for non-literal phenomena (e.g., Spotorno et al., 2012; van Ackeren et al., 2012; Feng et al., 2017), some individual study activation peaks and two ALE clusters had a high probability of falling within the ToM network. What is the role of mentalizing—a capacity supported by the ToM network— in language comprehension?

Because linguistic inputs often underspecify intended meaning (Wittgenstein, 1953; Sperber & Wilson, 1986), language comprehension routinely requires inferences about communicative intent. The computation of such inferences has historically been a focus of the field of pragmatics (Grice, 1957, 1975). Some early proposals drew a sharp boundary between literal and non-literal/pragmatic processing (Grice, 1975; Searle, 1979). However, defining the scope of pragmatic inference has proven challenging, and many have questioned the divide between literal and inferred meaning, or between semantics and pragmatics (Jackendoff, 2002). But if no such boundary exists, does language understanding *always* recruit the ToM network, in addition to the language network, to enable inferences about communicative intent?

The notion of continuous ToM engagement during language comprehension does not seem *a priori* plausible (many phenomena requiring context-based inferences—lexical disambiguation or pronoun resolution—are so common that it would seem inefficient and unnecessary to constantly engage in full-blown mentalizing) and does not find empirical support (Deen et al., 2015; Paunov et al., 2021; Shain, Paunov, Chen et al., 2022). Yet, in some cases, the ToM network does appear to contribute to language comprehension (e.g., Spotorno et al., 2012). Delineating the precise conditions under which comprehension requires ToM resources remains an important goal for future work (Paunov et al., 2021). Brain-imaging investigations of diverse non-literal phenomena using approaches with functionally localized ToM and language networks may offer some clarity. This approach could be complemented by experiments testing linguistic abilities in individuals with impaired ToM reasoning (e.g., Siegal et al., 1996; Happé et al., 1999) or in statistical language models (Devlin et al., 2019). The latter can reveal which non-literal phenomena can be handled by language models, and which might require additional machinery that approximates mental state inference (e.g., Hu et al., 2021).

Many individual study activation peaks and four ALE clusters had a high probability of falling within the language network. Of course, understanding both literal and non-literal utterances requires language processing, so both conditions should elicit a strong response in the language network. But why do non-literal conditions often elicit a stronger response? Differences in linguistic complexity might provide one explanation. The brain’s language areas are sensitive to comprehension difficulty (Wehbe et al., 2021). Although studies that compare literal and non-literal conditions commonly match stimuli on some linguistic variables, this matching is often limited to word-level features, which do not account for potential differences in context-based processing difficulty (e.g., surprisal; Smith & Levy, 2011). Further, despite the robust dissociation between the language and ToM networks, Paunov et al. (2019) found that the two networks show reliable correlation during language processing. This inter-network interaction may lead increased ToM demands to additionally manifest in the language system via inter-network connections. Finally, the language network may actually support some pragmatic computations, although to argue for such effects, linguistic confounds and inter-network information leakage would need to be eliminated as possible explanations.

### No evidence for the role of the MD network in non-literal comprehension

In contrast to the ToM and language networks, the Multiple Demand network does not appear to support non-literal comprehension: individual-study activation peaks and ALE clusters were least likely to fall in this network. Selecting the non-literal interpretation of an utterance may tax working memory (because multiple interpretations may be activated) or require inhibitory control (e.g., Gernsbacher & Robertson, 1999; Channon & Watts, 2003). However, recent work has shown that any such operation related to linguistic processing appears to be implemented within the language-selective system (Shain, Blank et al., 2020; see Fedorenko & Shain, 2021 for a review). Indeed, as discussed above, greater cognitive demands associated with non-literal interpretation may well explain the responses in the *language* system.

We suspect that neural activity observed during non-literal processing in putative executive control areas in past studies (e.g., AbdulSabur et al., 2014; Chan & Lavallee, 2015; Bosco et al., 2017) is due either to i) reliance on anatomical landmarks, which do not warrant functional interpretation in the association cortex (Poldrack, 2006; Fedorenko, 2021), or ii) extraneous task demands (see Diachek, Blank, Siegelman et al., 2020 for evidence that sentence comprehension only engages the MD network when accompanied by a secondary task, like a sentence judgment). Behavioral investigations that find correlations between executive abilities and non-literal interpretation abilities (e.g., Akbar et al., 2013; Caillies et al., 2014; Rints et al., 2015) are also likely affected by methodological issues, from small sample sizes to confounded experimental paradigms (Matthews et al., 2018). Indeed, recent large-scale investigations (Cardillo et al., 2020; Fairchild & Papafragou, 2021) argue against the role of executive functions in pragmatic ability.

### Possible reasons for the inconsistencies regarding the hemispheric bias

One remaining puzzle concerns the difference between patient work, which has implicated the RH in non-literal language processing (e.g., Winner & Gardner, 1977; Myers & Linebaugh, 1981; Delis et al., 1983; Brownell et al., 1983; 1986; Van Lancker & Kempler, 1987; Brownell, 1988; Stemmer et al., 1994; Giora et al., 2000; Ferstl et al., 2005) and our results, which demonstrate a LH bias. As noted above, left-lateralized activations have also been reported in past studies (e.g., Rapp et al., 2004; Lee & Dapretto, 2006; Hillert & BuracLas, 2009; Piñango et al., 2015; Bosco et al., 2017) and meta-analyses (Bohrn et al., 2012; Rapp et al., 2012; Reyes-Aguilar et al., 2018; Farkas et al., 2021). One possibility is that no RH bias exists for non-literal language processing. Both hemispheres contribute, perhaps with the LH contributing more strongly, as per the standard LH language bias (e.g., Geschwind, 1970). Because lexical and grammatical impairments resulting from LH damage are more salient and devastating, non-literal comprehension difficulties go unnoticed. In contrast, RH damage, which does not strongly affect basic language processing, may make apparent more subtle linguistic impairments, leading to the apparent RH bias for non-literal comprehension. In line with this possibility, several studies have reported that patients with LH damage show similar, or even greater, deficits in non-literal comprehension compared to RH-damaged patients (e.g., Tompkins, 1990; Giora et al., 2000; Zaidel et al., 2002; Gagnon et al., 2003; Klepousniotou & Baum, 2005; Ianni et al., 2014; Cardillo et al., 2018; Klooster et al., 2020).

Alternatively, the RH may indeed contribute more strongly than the LH to non-literal language comprehension, and past fMRI studies may have failed to detect the RH effects due the generally low sensitivity of the group-averaging analytic approach (Nieto-Castañón & Fedorenko, 2012). This possibility seems unlikely: if the RH were more active than the LH during non-literal comprehension, then RH activations should be easier to detect in a group analysis. This is because stronger responses would be more spatially extensive in individual brains and thus more likely to lead to overlap at the group level. (Further, using our probabilistic functional atlases, we did not find evidence for the possibility that functional areas in the RH are more variable in their locations across individuals and therefore less likely to emerge in a group analysis.) Future fMRI studies where areas of interest are identified via functional localizers, as well as intracranial stimulation studies, which enable spatially precise perturbations of neural activity, can provide important insight into the hemisphere-bias question.

### The need to carve non-literal processing at its joints

We did not find any systematic patterns with regard to the phenomena that ‘load’ onto the language vs. the ToM network (**SI Table 2**). However, many phenomena had small numbers of data points. In general, the field would benefit from clear, formal hypotheses about differences in the cognitive processes that contribute to each non-literal phenomenon. Such hypotheses can then be tested using behavioral, brain imaging, and computational modeling approaches.

We would like to conclude with a methodological point. fMRI has been a critical tool in human cognitive neuroscience. However, two disparate approaches are currently in use: i) the traditional approach where brains are averaged voxel-wise in the common space (Friston et al., 1999), and ii) the subject-specific functional localization approach where regions of interest are identified using functional ‘localizers’ and critical effects are examined therein (Saxe et al., 2006). Until recently, comparing findings across these approaches—and thus establishing a cumulative research enterprise—has proven challenging. Probabilistic functional atlases, like the ones we used in the current study (see also Dworetsky et al., 2021; Thirion et al., 2021), can bridge these two approaches by providing a common framework for functional areas/networks.

## Acknowledgements

We would like to thank Alex Paunov, Josef Affourtit, and Ben Lipkin for help with some of the analyses, and Ariel Goldberg and Jayden Ziegler for comments on the undergraduate thesis that resulted from this work. This work was supported by the NIH R01 award DC016607 and funds from MIT’s Simons Center for the Social Brain (via SFARI); EF was further supported by the NIH R01 award DC016950, and funds from the McGovern Institute for Brain Research and the Brain and Cognitive Sciences department

## Supplementary Information

**SI Table 1.**
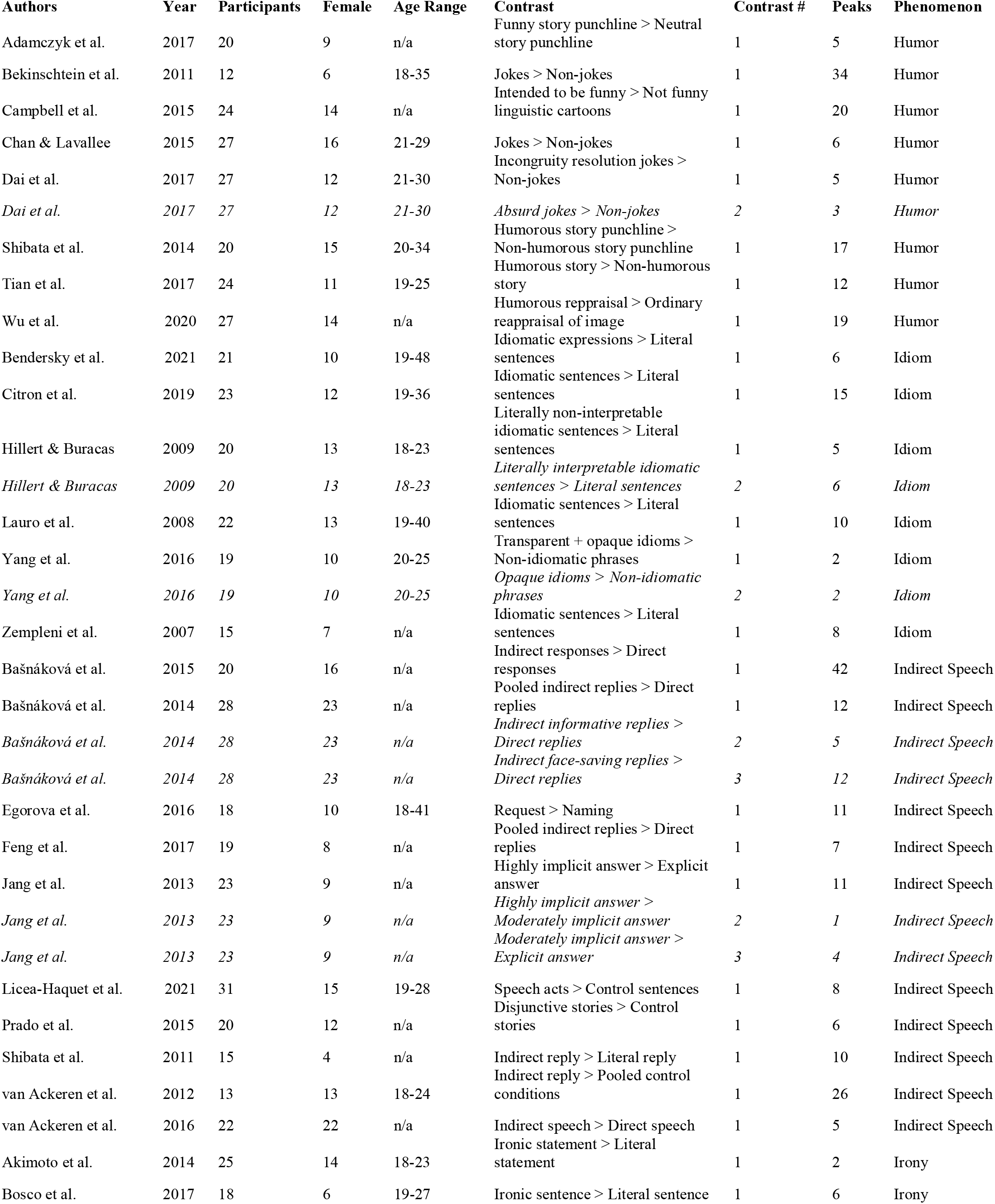
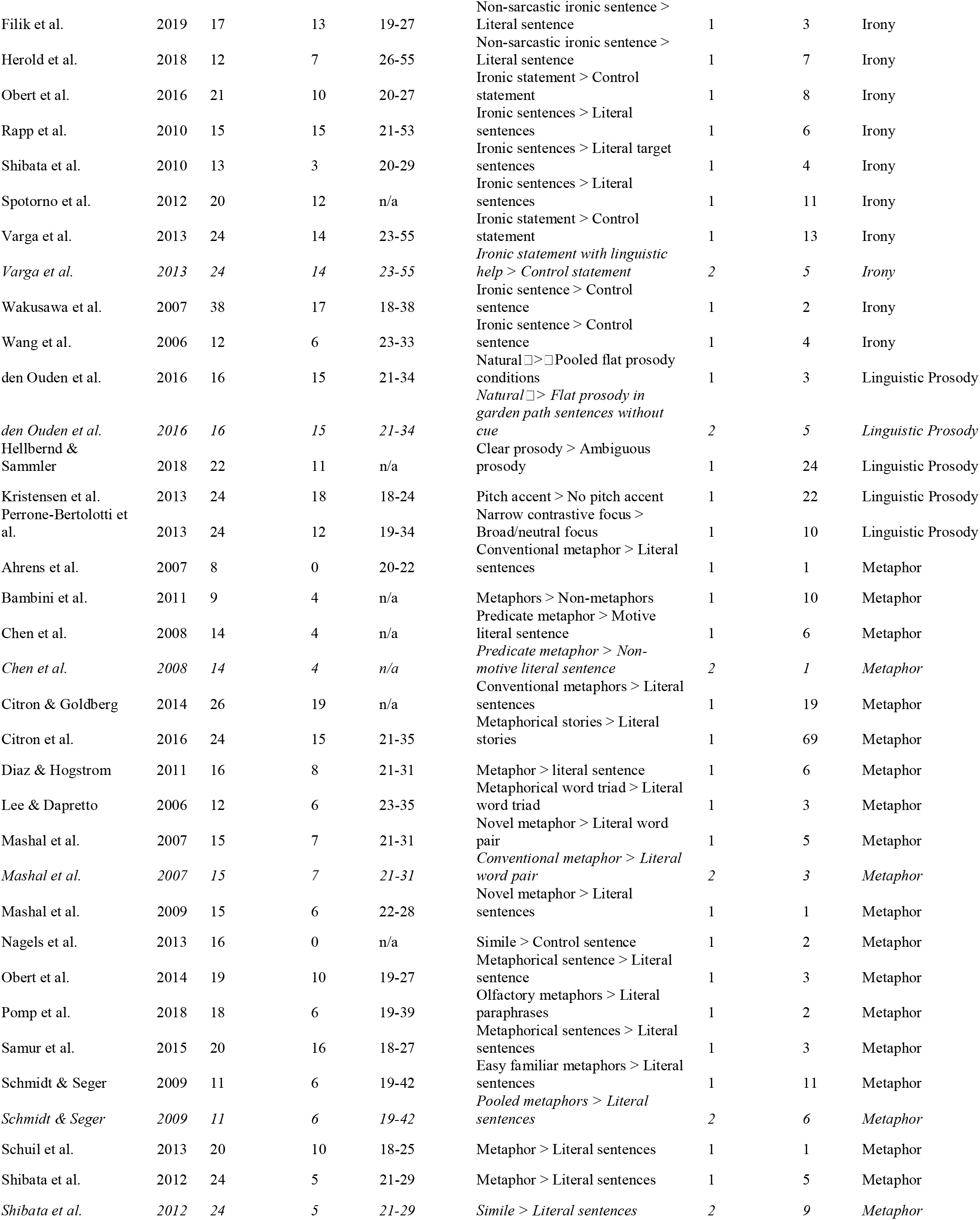
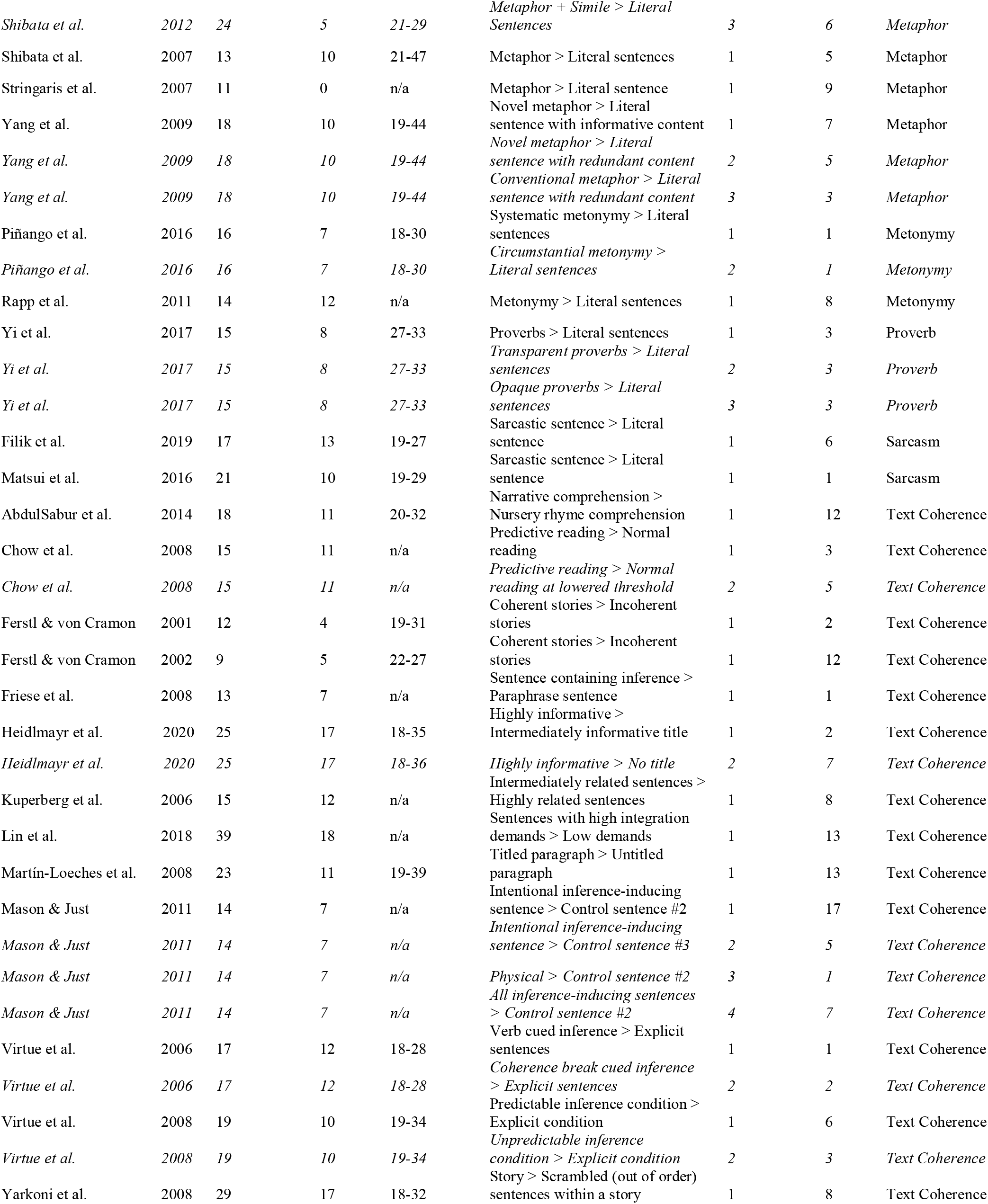
All individual contrasts included in the meta-analysis. Multiple contrasts taken from the same publication are displayed in italics.

**SI Table 2.** Individual peaks contributing to the six significant ALE clusters by phenomenon. Clusters (numbers the same as in Table 3) are listed by column. Empty space indicates that zero peaks from a given phenomenon contributed to the cluster assigned to that column.

|  | 1 | 2 | 3 | 4 | 5 | 6 |
| --- | --- | --- | --- | --- | --- | --- |
| **Humor** | 3 |  | 8 | 3 | 5 |  |
| **Idiom** | 1 |  | 5 | 2 | 1 | 1 |
| **Indirect Speech** | 3 | 4 | 20 |  | 7 | 11 |
| **Irony** | 5 | 4 | 8 |  | 2 | 3 |
| **Metaphor** | 1 |  | 19 | 2 | 6 | 6 |
| **Metonymy** |  | 1 | 1 |  |  | 1 |
| **Prosody** |  | 2 | 1 | 1 | 2 |  |
| **Proverb** |  | 2 | 1 |  |  |  |
| **Sarcasm** |  |  | 1 |  |  |  |
| **Text Coherence** | 2 | 4 | 17 | 2 | 5 | 1 |

**SI Figure 1.**
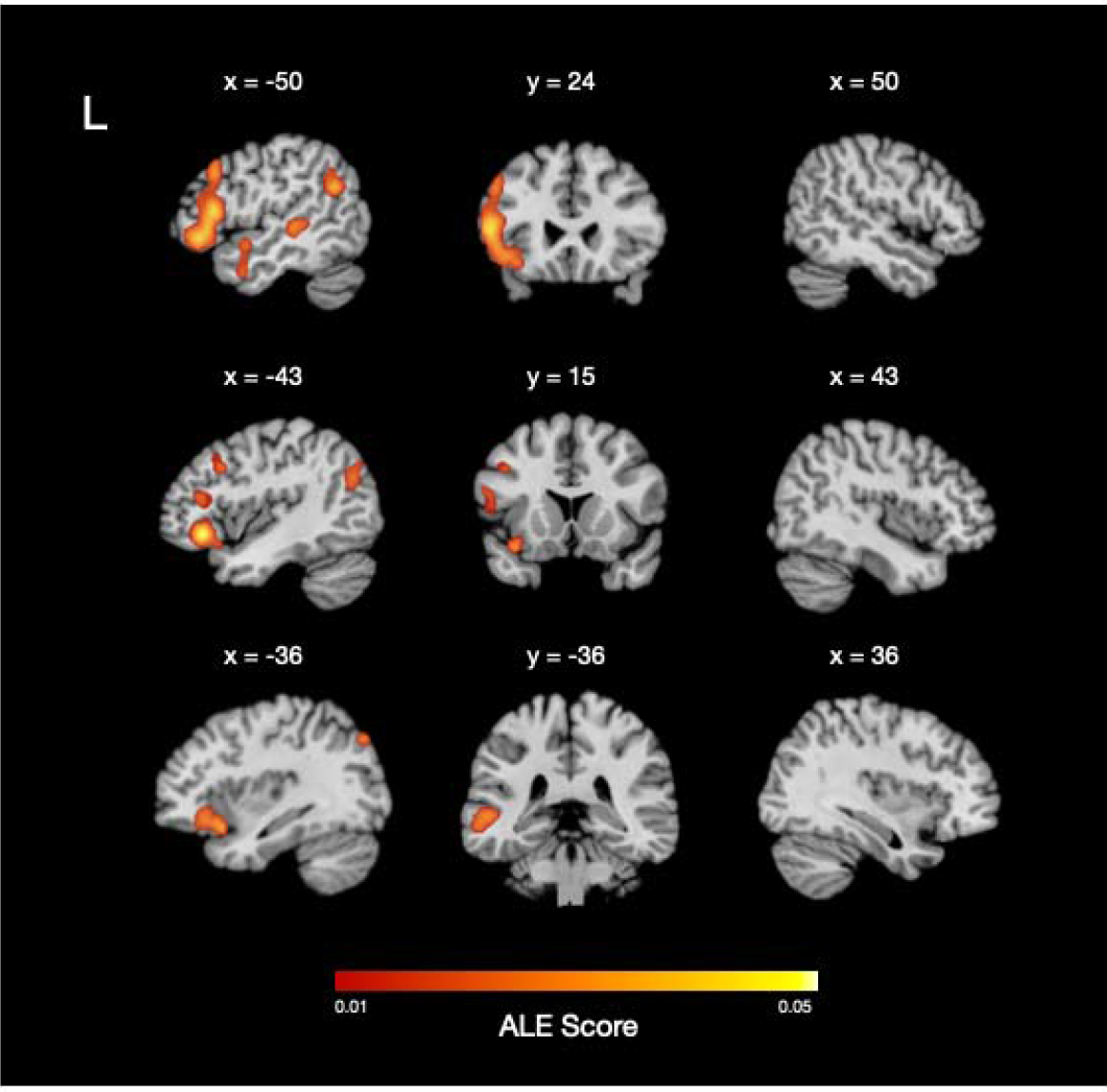
Main ALE results. ALE results for the 825 total peaks in our dataset, where peaks from each publication (n=74) are considered independent. ALE scores associated with each of the 6 significant ALE clusters are overlaid on a high-resolution structural MRI scan in MNI space (cluster-forming threshold = p<0.001 uncorrected, cluster-level family-wise error (FWE) = p<0.05, 1000 permutations).

